# Microbiome stability is linked to coral thermotolerance

**DOI:** 10.1101/2024.10.03.616497

**Authors:** Jake Ivan P. Baquiran, John Bennedick Quijano, Madeleine J.H. van Oppen, Patrick C. Cabaitan, Peter L. Harrison, Cecilia Conaco

**Affiliations:** Marine Science Institute, University of the Philippines Diliman, Quezon City, 1101, Philippines; Graduate School of Engineering and Science, University of the Ryukyus, 1 Senbaru, Nishihara, Okinawa 903-0213, Japan; Australian Institute of Marine Science, PMB No 3, Townsville MC, Queensland 4810, Australia; School of BioSciences, University of Melbourne, Parkville, Victoria 3010, Australia; Faculty of Science and Engineering, Southern Cross University, Lismore, NSW 2480, Australia

**Keywords:** thermotolerance, *Acropora* cf. *tenuis*, microbial community, *Endozoicomonas*, intraspecific variation, 16S rRNA sequencing, *Vibrio*

## Abstract

Corals associate with a diverse community of prokaryotic symbionts that provide nutrition, antioxidants, and other protective compounds to their host. However, the influence of microbes on coral thermotolerance remains understudied. Here, we examined the prokaryotic microbial communities associated with colonies of *Acropora* cf. *tenuis* that exhibit high or low thermotolerance upon exposure to 33°C (heated) relative to 29°C (control). Using 16S rRNA sequencing, we show that the microbial community structure of all *A.* cf. *tenuis* colonies were similar at control temperature. Thermotolerant colonies, however, had relatively greater abundance of *Endozoicomonas, Arcobacter*, *Bifidobacterium* and *Lactobacillus*. At elevated temperature, only thermosensitive colonies showed a distinct shift in their microbiome, with an increase in Flavobacteriales, Rhodobacteraceae, and *Vibrio,* accompanying a marked bleaching response. Functional prediction indicated that prokaryotic communities associated with thermotolerant corals were enriched for genes related to metabolism, while microbiomes of thermosensitive colonies were enriched for cell motility and antibiotic compound synthesis. These differences may contribute to the variable performance of thermotolerant and thermosensitive corals under thermal stress. Identification of microbial taxa correlated with thermotolerance provides insights into beneficial bacterial groups that could be used for microbiome engineering to support reef health in a changing climate.

## INTRODUCTION

Reef-building corals are essential components of coral reef ecosystems, laying the foundations of complex reefs and influencing biogeochemical processes. Corals are holobionts that host Symbiodiniaceae and prokaryotic microbial symbionts, as well as other microorganisms within and on their tissues (Thompson et al., 2014). These symbionts capture and recycle nutrients, provide food for their host, and produce antioxidants and protective compounds (Garrido et al., 2020; Matthews et al., 2020). Specifically, the bacterial members of the coral holobiont contribute to coral protection (e.g. antibiotic activity and predation of opportunistic pathogens) and metabolism (e.g. elemental cycling and synthesis of secondary metabolites) (Krediet et al., 2013; McDevitt-Irwin et al., 2017) to support the survival of corals in shallow oligotrophic waters.

Unprecedented rates of ocean warming and acidification threaten marine ecosystems and will impact survival of vulnerable marine species (Veron et al., 2009; Doney et al., 2020; Giddens et al., 2022). Rising seawater temperature, in particular, is a major factor that can disrupt mutualistic interactions within the coral holobiont, resulting in bleaching and mortality (Miller et al., 2009; Rädecker et al., 2020). Bleaching refers to the loss of the microalgal endosymbionts and/or their photopigments from the coral tissues (Hoegh-Guldberg, 1999; Takahashi et al., 2008; Oakley and David, 2018; Helgoe et al., 2024). Elevated temperature has been correlated with the spread of coral diseases as a warming ocean may facilitate the proliferation and promote virulence of potential microbial pathogens (Miller and Richardson, 2014). When bleaching occurs, the immunity of the coral host is compromised, resulting in changes in coral-associated bacteria and an increase in susceptibility to diseases (Muller et al., 2018; Kusdianto et al., 2021; Cavanich et al., 2022). In recent decades, many instances of mass coral bleaching during summer heatwaves have been reported in Atlantic, Caribbean, and Indo-Pacific coral reefs (Neal et al., 2017; Lough et al., 2018; Sully et al., 2019; Quimpo et al., 2020; Pereira et al., 2022; Henley et al., 2024). These events are predicted to increase in frequency and severity and are the major driver of continued degradation of coral reefs worldwide.

Observations of coral communities on reefs experiencing mass bleaching events have revealed intraspecific variation in bleaching susceptibility (Penin et al., 2007; Morikawa and Palumbi, 2019; Chapron et al., 2022). When bleaching occurs, some individuals will die while others within the same population are able to recover; hence some colonies are more resistant to thermally-induced bleaching. Although thermotolerance is often attributed to the microalgal associates of corals (Oliver and Palumbi, 2011; Palacio-Castro et al., 2023), several studies have demonstrated that the taxonomic composition of Symbiodiniaceae does not always correlate with observed heat tolerance of corals (Humanes et al., 2021, 2022). Additionally, prokaryotic microbial community dynamics has been linked to coral thermotolerance, with resistant colonies often exhibiting more stable microbiomes compared to susceptible colonies (Palacio-Castro et al., 2022). For instance, in *Acropora hyacinthus* from different reef sites, heat-sensitive individuals observed at a site with a history of moderate temperature fluctuations exhibited changes in their microbiome, while heat-tolerant individuals from a site experiencing high temperature extremes remained unchanged (Ziegler et al., 2017). In addition, metagenomes of mucus-associated microbes from resilient individuals of the coral *Pseudodiploria strigosa*, sampled from both inner and outer reefs exposed to heat stress, showed greater similarity at the level of bacterial genera with an increase in abundance of beneficial taxa, as well as genes related to stress response and metabolism of nitrogen and sulfur when compared to ambient treatments (Lima et al., 2024). However, whether prokaryotic community differences underlie variability in thermal tolerance of corals within the same reef habitat remains to be investigated.

Here, we compared the prokaryotic community associated with thermotolerant and thermosensitive colonies of *Acropora* cf. *tenuis* located within the same reef area in Northern Luzon, Philippines. We show that, although high and low tolerance corals had similar microbiome structure under ambient conditions, the microbiomes of susceptible colonies were more affected by thermal stress. We also identified microbial taxa that may be indicators of better performance at elevated temperature.

## MATERIALS AND METHODS

### Thermal stress exposure

Thirty healthy colonies of *Acropora* cf. *tenuis* from 4-5 m depth at Caniogan reef (16.29473 N, 120.01448 E) in Anda, Pangasinan, northwestern Philippines, were selected and tagged. Pieces of the coral colonies were collected with permission from the Philippines Department of Agriculture Bureau of Fisheries and Aquatic Resources (DA-BFAR Gratuitous Permit No. GP-0169-19). The coral pieces were transported to the outdoor hatchery of the Bolinao Marine Laboratory on November 13, 2021 where they were acclimated in tanks with flowing seawater at ∼29°C for a day. Sixteen fragments (∼1 cm in length) were then broken off from each of the colonies and attached to cement nubbin holders using cyanoacrylate glue (Dizon et al., 2008). Nubbins were labeled with small plastic tags to enable tracking of parent colonies.

Coral fragments attached to the holders were distributed into eight 40-L polycarbonate tanks with aerated and flow-through (ca. 20L/hr) sand-filtered seawater kept at 29°C. After 3 days of acclimation, the temperature in four tanks was gradually increased to 33°C (+1°C/day) to approximate the maximum summer seawater temperature in the Bolinao-Anda Reef Complex, while the other four tanks were maintained at 29°C (controls). Upon reaching the treatment temperature, conditions were maintained for another four days (Supplementary Figure 1a; Supplementary Table 1). Temperature was regulated using submersible heaters (Eheim, Germany) controlled by thermostats and additional water circulation was provided by 200L/hr submersible pumps (Atlantis HX-1000) for even distribution of heat. Lighting was provided by cool daylight LEDs (ca. 100 uE m-2 s-1). During the experiment, temperature and pH were monitored using a portable Vernier LabQuest 2 (Vernier, USA) and salinity and dissolved oxygen (DO) using a Horiba LAQUAact-PC110 multiparameter meter (HORIBA Advanced Techno, Co., Ltd., Japan).

After four days of sustained exposure to the experimental temperatures, coral colonies were classified into tolerance categories: (1) thermotolerant (if no mortality or signs of bleaching were observed in both 33°C and control tanks, (2) thermosensitive if ≥ 50% of the fragments from the colony showed bleaching in 33°C tanks but not in control tanks, and (3) unclassified if bleaching or mortality were observed during acclimation or in ambient tanks during the experiment.

### DNA extraction and sequencing

Two fragments were sampled from each of four colonies representing thermotolerant and thermosensitive individuals from the 29°C (hereinafter referred to as “T29” and “S29”, respectively) and 33°C (hereinafter referred to as “T33” and “S33”, respectively) treatments. All fragments were apparently healthy, except for representatives of S33, which showed bleaching and some tissue sloughing. Coral fragments were flash-frozen in liquid nitrogen and stored at −80°C prior to DNA extraction. Entire fragments were crushed using mortar and pestle, and total genomic DNA was extracted using a modified cetyltrimethylammonium bromide (CTAB) method (Winnepenninckx et al., 1993). DNA was resuspended in nuclease-free water and stored at −20°C. The quality of extracted DNA was checked by agarose gel electrophoresis using 1% agarose in 1X Tris/Borate/EDTA buffer. DNA concentration was determined using a NanoDrop 2000c spectrophotometer (Thermo Scientific, Waltham, MA, USA). Genomic DNA from fragments derived from the same colony and treatment were pooled together resulting in 16 samples representing 4 colonies per tolerance category and treatment. High quality total genomic DNA was submitted to Macrogen, Inc., South Korea, for 16S rRNA gene sequencing. Prokaryotic 16S rRNA gene (V3V4 hypervariable region) amplicon sequence libraries were prepared using Herculase II Fusion DNA Polymerase and the Nextera XT Index V2 Kit (Agilent Technologies, Santa Clara, CA, USA) with primers Bakt_341F (5’-CCTACGGGNGGCWGCAG-3’) and Bakt_805R (5’- GACTACHVGGGTATCTAATCC-3’) (Herlemann et al., 2011). Paired-end sequencing (300 bp) was performed on the MiSeq platform (Illumina, Inc., San Diego, CA, USA) following the dual-index sequencing strategy.

### Microbial community analysis

Microbial community analysis was conducted using the Quantitative Insights Into Microbial Ecology 2 (QIIME2) package (2021.11) (Bolyen et al., 2019; https://docs.qiime2.org/2021.11/). Raw sequence data were deposited in the NCBI Sequence Read Archive and can be accessed under BioProject accession number PRJNA1033317. Raw data are also available on Figshare (https://doi.org/10.6084/m9.figshare.24715299). Primers were trimmed from the reads using cutadapt. Reads were merged into amplicon sequence variants (ASVs) using the DADA2 package (Callahan et al., 2016) following these parameters: -p-trunc-len-f 260 –p-trunc-len-r 200. Singleton and doubleton ASVs were removed prior to annotation. SILVA version 138.1 (Quast et al., 2012; https://arb-silva.de) was used for taxonomic classification. Mitochondrial and chloroplast sequences were removed from the final set of ASVs.

Rarefaction curves were generated based on Pielou’s evenness and Shannon diversity. Alpha diversity metrics, including Faith’s phylogenetic diversity (PD), Shannon, observed features, and Chao1 were calculated from microbiome data rarefied to the smallest sequence library. Comparison of alpha diversity among groups was done using Kruskal-Wallis test. Results of pairwise Kruskal-Wallis tests were adjusted using Benjamini-Hochberg (BH) false discovery rate correction procedure (Benjamini and Hochberg, 1995). Community distance matrices based on Bray-Curtis dissimilarity were visualized using Principal Coordinates Analysis (PCoA). PERMANOVA using the adonis2 function with 999 permutations was used to test for associations between microbiome composition and heat tolerance classifications or temperature treatments. Differentially abundant ASVs (BH-adjusted p-value <0.05) were identified using Analysis of Compositions of Microbiomes with Bias Correction (ANCOMBC2) (Lin and Peddada, 2020, 2024).

### Endozoicomonas phylogeny

Near full-length *Endozoicomonas* sequences were extracted from RefSeq (Pruitt, 2004), aligned using MAFFT, and trimmed using ClipKit (Steenwyk et al., 2020). Reference trees were constructed using maximum likelihood (ML) on IQ-TREE 2.0 (Katoh and Standley, 2013; Minh et al., 2020). Branch support was calculated with 1,000 ultrafast bootstrap replicates (UFBS; ‘-B 1000’ command) and Shimodaira–Hasegawa approximate likelihood ratio tests (SH-aLRT; ‘-alrt 1000’ command) (Ota et al., 2000; Minh et al., 2013). *Endozoicomonas* ASVs were extracted from our sequence data and their probable position on the reference tree was estimated by maximum likelihood using EPA-NG (Barbera et al., 2018).

### Functional prediction

Functional gene abundance based on ASV taxon affiliations was predicted using Phylogenetic Investigation of Communities by Reconstruction of Unobserved States or PICRUSt2 (Langille et al., 2013). Differentially abundant KEGG orthologs (KO) were identified using ANCOMBC2 (Lin and Peddada, 2020, 2024). Differentially abundant KOs (q-value <0.05) with total relative abundance ≥ 0.02% were grouped into functions based on KEGG BRITE hierarchy level 2 for visualization.

### Data visualization

Data processing and visualization were done using tidyverse (Wickham et al., 2019), a collection of R packages (R Core Team, 2021), implemented in RStudio (RStudio Team, 2020).

## RESULTS

### Microbial community structure of Acropora cf. tenuis

Exposure of *A.* cf. *tenuis* colonies to control (29°C) or elevated (33°C) temperature treatments elicited variable responses that indicate inter-individual differences in thermotolerance (Figure 1). While thermosensitive colonies exhibited bleaching and some tissue sloughing at 33°C, thermotolerant colonies remained apparently healthy with no signs of bleaching even after 4 days of sustained exposure to high temperature (Supplementary Figure 1b; Supplementary Table 2).

**Figure 1.**
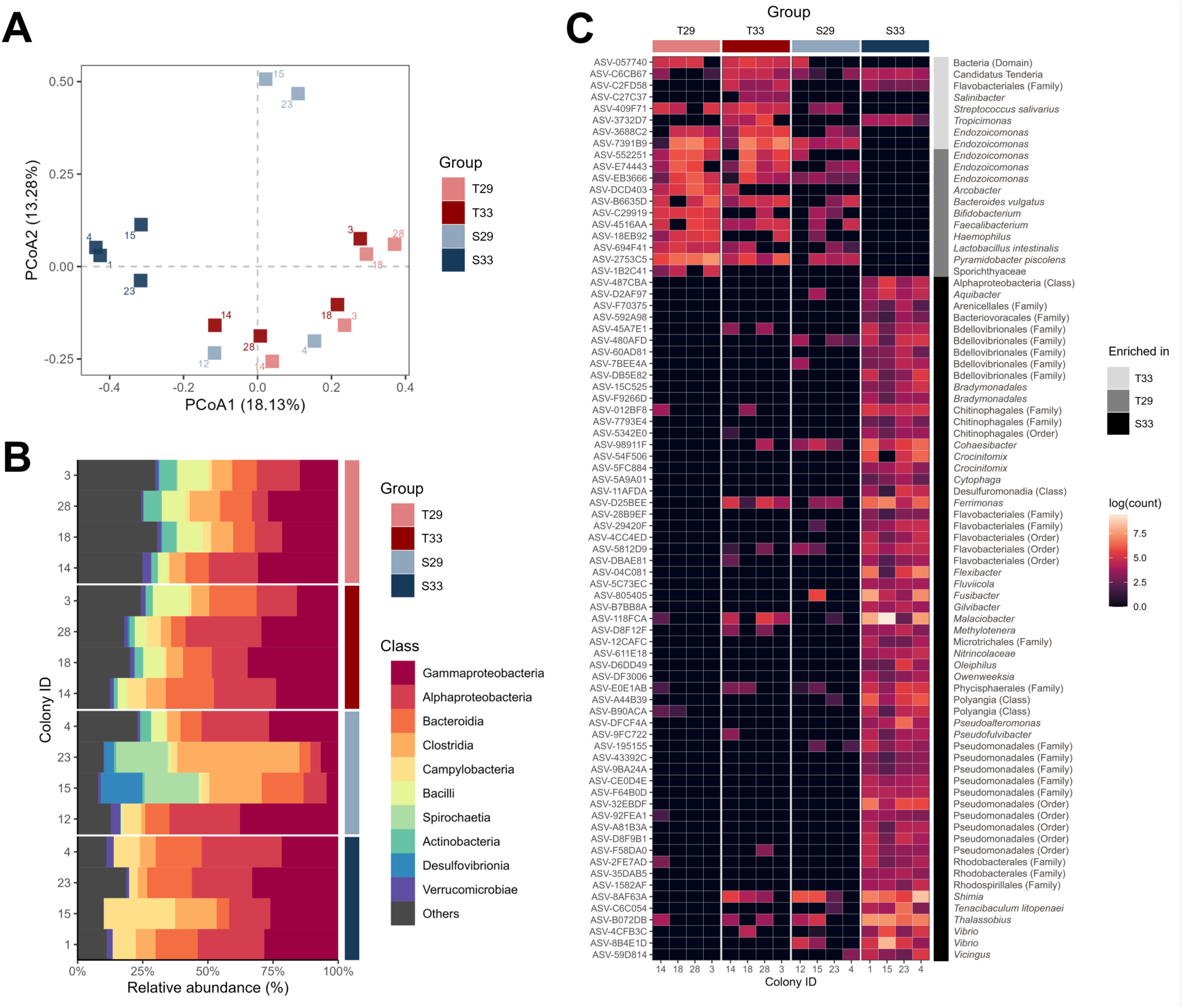
Composition and structure of microbial communities associated with *A.* cf*. tenuis.* (a) Principal coordinates analysis (PCoA) of microbial communities of thermotolerant (T) or thermosensitive (S) colonies subjected to 33°C or 29°C treatments (T29, light red; T33, dark red; S29, light blue; S33, dark blue). Colony IDs are shown beside the markers. (b) Relative abundance of the top 10 microbial classes in the *A.* cf*. tenuis* microbiome. (c) Heatmap of differentially abundant ASVs (p-value < 0.05). Heatmap colors represent log-normalized counts.

To examine whether observed differences in bleaching response correlated with the prokaryotic associates of the colonies, we characterized the microbial community of selected samples. Sequencing of the 16S rRNA gene (V3V4 region) from 16 libraries representing thermotolerant and thermosensitive colonies that had been subjected to control or elevated temperature treatments (n = 4) yielded a total of 1,474,619 raw reads. From these, 1,295,805 high-quality reads were retained after sequence filtering, with an average of 80,987.82 ± 9,178.42 (mean ± standard deviation) reads per library. Further filtering generated 5,718 unique amplicon sequence variants (ASVs) with a total frequency of 826,847 at an average of 51,677.94 ± 9,258.04 (mean ± standard deviation) per library. Rarefaction curves based on Pielou’s evenness and Shannon diversity reached a plateau at around 5,000 sequences, indicating that sequencing depth even for the smallest library (with 35,588 sequences) provided sufficient coverage of the prokaryotic communities in our samples (Supplementary Figure 2). ASVs associated with the corals were classified into 48 phyla, 113 classes, 273 orders, 466 families, and 872 genera.

No significant differences in alpha diversity were observed among thermotolerant and thermosensitive corals based on Faith’s PD, Shannon, observed ASVs, and Chao1 indices (Supplementary Figure 3). The *A.* cf. *tenuis*-associated prokaryotic microbiome was dominated by the phyla Proteobacteria (40.64%), Firmicutes (16.72%), Bacteroidota (12.22%), Campylobacterota (5.57%), Actinobacteriota (3.39%) and the classes Gammaproteobacteria (22.62%), Alphaproteobacteria (18.02%), Bacteroidia (11.6%), Clostridia (11.45%), Campylobacteria (5.57%) (Figure 1b, Supplementary Figure 4).

The core *A.* cf. *tenuis* microbiome comprised 2.41% (138 ASVs) of ASVs that were shared among all treatment groups (Supplementary Figure 5). Regardless of temperature treatment, 96 and 98 ASVs were present in all thermotolerant and thermosensitive colonies, respectively. Conversely, 69 and 140 ASVs were present in all colonies maintained at ambient or elevated temperature, respectively, regardless of tolerance category. Unique ASVs comprised 16.27% to 23.21% of the coral-associated microbial community of each group of samples.

### Influence of tolerance class and temperature on the coral microbiome

Microbial community structure was influenced by tolerance class and temperature treatment (Figure 1c; Data Files 1-2). Principal coordinate analysis revealed clustering of all thermotolerant samples, together with some thermosensitive samples from the ambient temperature treatment, whereas all thermosensitive samples from the heated treatment formed an independent cluster (Figure 1a).

At ambient temperature, microbial communities in thermotolerant and thermosensitive colonies showed no significant difference in structure (PERMANOVA, p-value=0.171; Table 1). Only 6 ASVs were found at significantly greater abundance in thermotolerant relative to thermosensitive corals based on ANCOMBC2. These ASVs were affiliated with *Arcobacter* (Campylobacteraceae), *Haemophilus* (Pasteurellaceae), *Bifidobacterium* (Bifidobacteraceae), *Lactobacillus* (Lactobacillaceae), *Pyramidobacter* (Synergistaceae), and Sporichthyaceae (Figure 1c, Data Files 1-2).

**Table 1.** Comparison of *Acropora* cf*. tenuis* microbial communities at ASV level between heat tolerance classifications and temperature treatments based on Bray-Curtis dissimilarity. Benjamini-Hochberg adjusted p-values in bold denote statistical significance at p < 0.05.\

|  | **PERMANOVA** |  |
| --- | --- | --- |
| **Global comparisons** | **pseudo-F** | **p-value** |
| Tolerance classification | 1.933095 | **0.002** |
| Temperature treatment | 2.135664 | **0.048** |
| Tolerance x Treatment | 1.736775 | **0.002** |
| **Pairwise comparisons** |  |  |
| T29 x T33 | 0.955802 | 0.550 |
| T29 x S29 | 1.224929 | 0.171 |
| T29 x S33 | 2.929221 | **0.034** |
| T33 x S29 | 1.158406 | 0.356 |
| T33 x S33 | 2.408634 | **0.036** |
| S29 x S33 | 2.341153 | **0.029** |

At elevated temperature, the microbiome structure of thermotolerant and thermosensitive colonies behaved differently (PERMANOVA, p-value=0.036; Table 1). Thermotolerant microbiomes remained similar to controls at ambient temperature (PERMANOVA, p-value=0.55; Table 1), with only 5 ASVs (*Salinibacter*, Candidatus *Tenderia*, *Tropicimonas*, *Ferrimonas,* and Cryomorphaceae) showing significant enrichment, and 4 ASVs (Sporichthyaceae, *Haemophilus*, *Arcobacter*, and *Bifidobacterium*) showing depletion (Figure 1c, Data File 2). In contrast, thermosensitive colonies in the high temperature treatment exhibited a significant shift in microbial structure relative to controls (PERMANOVA p-value=0.029; Fig. 1a; Table 1; Supplementary Figure 4; Supplementary Table 4). Thermosensitive colonies at 33°C were enriched for 49 ASVs, including 17 Bacteroidia, 16 Gammaproteobacteria (12 Pseudomonadales, 2 Vibrionaceae), 4 Alphaproteobacteria (2 Rhodobacteraceae, 1 Terasakiellaceae, 1 unclassified), 4 Bdellovibrionaceae, 3 Desulfuromonadia, 1 Acidomicrobiia (Ilumatobacteraceae), 1 Campylobacteria (Malaciobacter), 1 Phycisphaeraceae, and 2 Polyangia, whereas 2 ASVs ( Endozoicomonaceae (*Endozoicomonas*) and Synergistaceae (*Pyramidobacter*) were significantly depleted (Figure 1c; Supplementary Table 4; Data File 2).

### Endozoicomonas are enriched in high tolerance colonies

We identified 28 ASVs affiliated with *Endozoicomonas* in the *A.* cf. *tenuis* microbiome (Figure 2). 14 ASVs were affiliated with *Endozoicomonas* sp. P9H01 isolated from coral, 1 ASV with *E. gorgoniicola*, 8 ASVs with *E. euniceicola*, and 5 ASVs with *Parendozoicomonas haliclonae* (Supplementary Figure 6). *Endozoicomonas* ASVs comprised 1.53% of the thermotolerant microbiome and only 0.23% of the thermosensitive microbiome under ambient temperature. Only 3 *Endozoicomonas* ASVs (0.0051%) were detected in thermosensitive colonies at 33°C (Data File 1). Five *Endozoicomonas* ASVs, including the 2 most abundant sequences, showed significant depletion in thermosensitive relative to thermotolerant colonies in the heated treatments.

**Figure 2.**
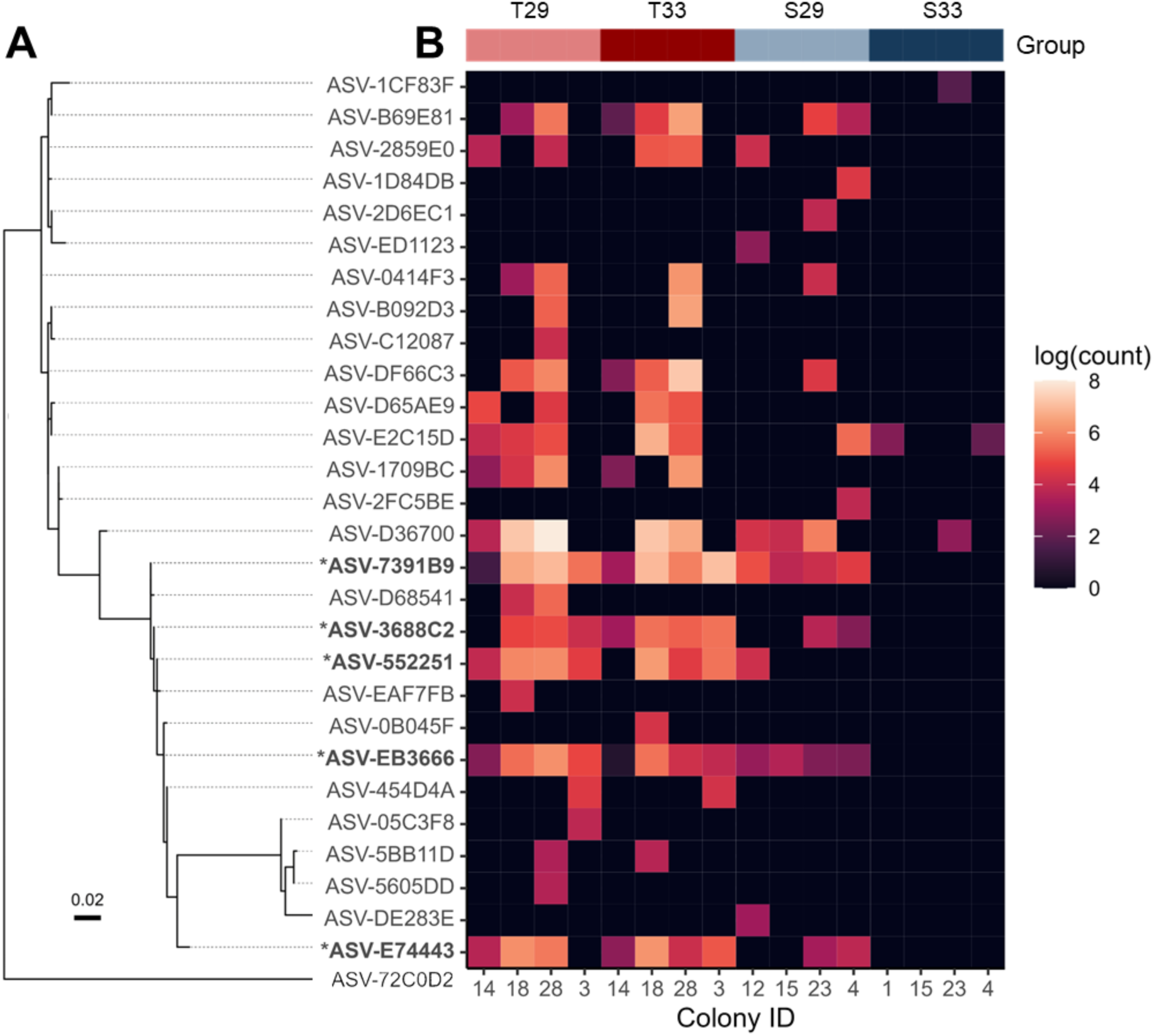
*Endozoicomonas* ASVs in the microbiome of *A.* cf. *tenuis*. (A) Maximum-likelihood tree of *Endozoicomonas* ASVs. The tree is rooted on *Neptunibacter* sp. (ASV-72C0D2). Asterisks indicate differentially abundant ASVs. (B) Heatmap of log-normalized counts of ASVs across thermotolerance categories and treatments (T29, light red; T33, dark red; S29, light blue; S33, dark blue).

### Predicted functions overrepresented in coral microbiomes

Functions associated with the microbiomes of *A. cf. tenuis* were predicted using PICRUSt2 (Figure 3). A total of 7,731 KEGG ortholog (KO) genes were represented in the microbiomes of *A.* cf. *tenuis* corals. The predicted KO profiles showed no clear differentiation between tolerance classes (PERMANOVA: R^2^ = 0.1, p-value = 0.195; Supplementary Figure 7). Thermotolerant corals showed relatively greater abundance of genes involved in metabolism of carbohydrates, lipids, and amino acids, and translation, compared to thermosensitive corals. In contrast, thermosensitive corals were enriched for genes involved in cell motility. When subjected to elevated temperature, the microbiome of thermotolerant corals remained similar to the microbiome at ambient temperature, whereas in the thermosensitive corals, functions related to carbohydrate, lipid, and amino acid metabolism were depleted and terpenoid and polyketide metabolism was enriched.

**Figure 3.**
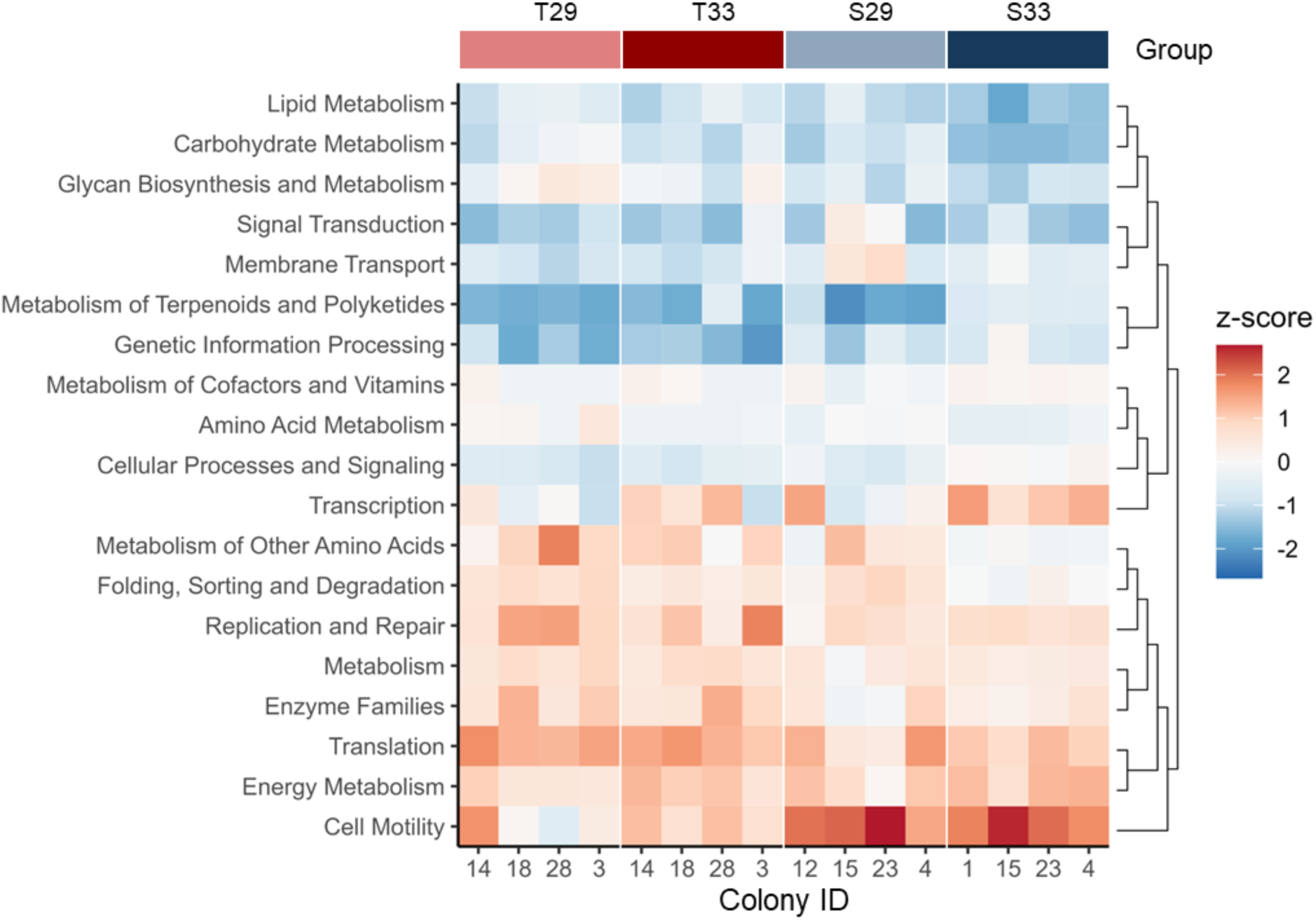
Differentially abundant predicted gene functions in the *A.* cf. *tenuis* microbiome. Only KEGG orthologs (KO) with total relative abundance ≥ 0.02% and q-value < 0.05 are represented. KOs were grouped into functions based on KEGG BRITE hierarchy level 2. Heatmap colors represent z-scores (red, high; blue, low).

## DISCUSSION

### Indicators of thermotolerance in the microbiome of A. cf. tenuis

Sympatric colonies of *A.* cf. *tenuis* exhibited variable temperature tolerance and could be grouped into thermotolerant and thermosensitive classes based on their bleaching responses upon exposure to elevated temperature. The native microbial community associated with colonies from the thermotolerant and thermosensitive groups did not show significant differences in structure under ambient conditions. However, distinct microbiome responses were observed when the corals were subjected to elevated temperature. Under these conditions, the microbiomes of thermotolerant corals remained comparable to controls, indicating community stability. On the other hand, the microbiomes of thermosensitive corals showed a significant change in community structure, becoming more homogenous in terms of composition. This is likely due to exclusion of heat-susceptible microbes and increased dominance of certain taxa (Sweet et al., 2019). Given the importance of coral-associated bacteria on holobiont performance (Voolstra et al., 2024), microbial community structure and measured responses to external stimuli may offer a useful indicator of coral health and resistance to stressors like elevated temperature (Palacio-Castro et al., 2022; Kusdianto et al., 2021; Chavanich et al., 2022).

A detailed analysis of the microbiomes of *A.* cf. *tenuis* highlighted specific microbial taxa that may relate to thermal tolerance characteristics. *Acropora* cf. *tenuis* is dominated by members of Gammaproteobacteria and Alphaproteobacteria, which concurs with findings by other groups that the microbiota of *A. tenuis* colonies is typically comprised of proteobacterial members (Quigley et al., 2019; Marchioro et al., 2020). The microbiome of thermotolerant colonies revealed greater abundance of *Arcobacter, Bifidobacterium*, *Lactobacillus,* and Sporichthyaceae compared to thermosensitive colonies. This suggests that members of these microbial groups may be positive predictors of coral thermotolerance. *Arcobacter* has been suggested to be associated with increased protein content reserves potentially benefiting the growth and development of *Pocillopora damicornis* (Zhang et al., 2021). Species belonging to *Bifidobacterium* are known members of the *Porites lutea* core microbiome (Gong et al., 2020) and have been reported to exert beneficial effects on host physiology and pathology including protection against lethal infections by pathogenic microbes (Fukuda et al., 2011). *Lactobacillus* dominated the *A. cervicornis* microbiome collected from deeper waters (Godoy-Vitorino, 2017) and is posited to exhibit antimicrobial activity against deleterious invader microbes (Chen et al., 2021). Sporichthyaceae, which are known to be facultative anaerobes, have been found in microbiomes impacted by wastewater suggesting that they can withstand stressful conditions (Newton and McLellan, 2015; Zárate et al., 2020).

Upon exposure of thermotolerant colonies to elevated temperature, changes in the relative abundance of several microbial taxa were noted. Taxa that increased in abundance include *Tropicimonas, Ferrimonas*, and Cryomorphaceae. *Tropicimonas* was previously identified as a bioindicator of microbiome recovery after bleaching in *Montipora peltiformis* (Zhang et al., 2023). *Ferrimonas* and Cryomorphaceae, in contrast, are potential microbial pathogens or opportunists and have been reported in the microbiome of diseased *Acropora palmata* (Rosales et al., 2019; Young et al., 2023). Taxa that decreased in abundance in thermotolerant colonies subjected to heat stress include *Haemophilus*, Sporichthyaceae, *Arcobacter* and *Bifidobacterium*, suggesting that these taxa are sensitive to temperature. *Haemophilus* has previously been reported to be less abundant in *Pocillopora damicornis* inoculated with beneficial actinobacterial strain (SCSIO 13291) exposed to heat stress (Li et al., 2023).

Specific members of the microbial assemblage of thermosensitive corals showed significant shifts in abundance accompanying the bleaching response at elevated temperature. Under these conditions, heat-susceptible microbes such as Bacilli, Desulfovibrionia, Actinobacteria and the archaea Halobacteria decreased in abundance. This depletion may be attributed to population elimination due to bacterial susceptibility to higher temperature or to competition by other microbiome members. Members of these prokaryotic taxa have previously been reported to be depleted in bleached coral fragments and in corals subjected to warming and acidification conditions (Sun et al., 2022; Grotolli et al., 2018).

Notably, many other microbial taxa in thermosensitive colonies increased in abundance during heat-induced bleaching, including members of Flavobacteria, Rhodobacteraceae, *Malaciobacter, Vibrio*, and Pseudomonadales. These groups of copiotrophic bacteria may proliferate in stressed corals due to the overproduction and shedding of carbon-rich mucus from the coral or exudates from degraded Symbiodiniaceae (Fujise et al., 2014; Tout et al., 2014; Lee et al., 2016). Flavobacteriales have been found in corals under stressful conditions and they are known to be initial members of the microbiome associated with stony coral tissue loss disease (McDevitt-Irwin et al., 2017; Rosales et al., 2023). Some Rhodobacteraceae are opportunist microbes that can compromise host health and are known as robust indicator species for thermally stressed corals (Pootakham et al., 2019; Hsieh et al., 2023). *Malaciobacter*, an emerging pathogenic genus for marine hosts, has been found at high abundance in *Drupella*-grazed *Acropora* corals and even in diseased marine bivalves (Auguste et al., 2022; Nguyen et al., 2023). *Vibrio* also includes pathogenic members that have been detected in diseased coral tissues and in stressed microbiomes of other acroporid corals (Gajigan et al., 2017; Meyer et al., 2019; Meenatchi et al., 2020; Schul et al., 2023). Metabolites released by certain *Vibrio* species favor the growth of co-pathogens, which can result in tissue necrosis (Rubio-Portillo et al., 2020). Interestingly, Bdellovibrionaceae, which are known predators of *Vibrios*, were also enriched in thermosensitive colonies under elevated temperature (Welsh et al., 2015). Finally, the increased abundance of Pseudomonadales in thermosensitive corals may be a mechanism to mitigate the impacts of high temperature, as members of this group are known to exert beneficial effects on corals through their antagonistic activity against disease-causing bacteria (Tiirola et al., 2022; Gignoux-Wolfsohn and Vollmer, 2015; Sun et al., 2022).

### Endozoicomonas abundance correlates with thermotolerance

*Endozoicomonas* comprised a substantial group within the *A.* cf. *tenuis* microbiome. Diverse *Endozoicomonas* ASVs were detected in thermotolerant *A.* cf. *tenuis* colonies at both ambient and elevated temperature conditions, with fewer in the thermosensitive group, especially in colonies subjected to high temperature. These results suggest that diversity and abundance of *Endozoicomonas* within the *A.* cf. *tenuis* microbial community may be an indicator of thermotolerance. Our findings mirror reports showing that *Endozoicomonas* are depleted in stressed, diseased, and bleached corals (Tandon et al., 2020; Botté et al., 2022). *Endozoicomonas* are known coral symbionts, contributing beneficial functions such as dimethylsulfoniopropionate (DMSP) transformation, amino acid and carbohydrate metabolism, and nutrient cycling (Tandon et al., 2019; Sweet et al., 2020; Pogoreutz et al., 2022). These bacteria can also convert cholesterol derived from the coral host into steroids that may inhibit pathogenic bacteria and fungi (Ochsenkühn et al., 2023). The abundance of *Endozoicomonas* may be considered as a gauge of coral health and may reflect coral recovery after a severe bleaching event (Chuang et al., 2024). However, other studies suggest that certain *Endozoicomonas* taxa may act as opportunistic microbes, switching between parasitism and pathogenicity depending on holobiont status (Pogoreutz and Ziegler, 2024). Further studies are therefore warranted to determine the exact functional roles and contributions of this group to coral thermotolerance.

### Microbiome functions mirror coral microbial community dynamics

Shifts in coral microbial community composition predict changes in the functions of the microbiome (Reigel et al., 2021; Bergman et al., 2023). The maintenance of metabolic functions for lipids, carbohydrates, glycans, and amino acids in the microbiome of thermotolerant *A.* cf. *tenuis* even under thermal stress may help maintain homeostasis, contributing to holobiont heat resistance (Zhang et al., 2015; Ziegler et al., 2017). In contrast, decreased abundance of metabolism-related genes in thermosensitive corals indicates weaker regulation of energy reserves and lowered protection against free radicals (Su et al., 2021). Moreover, enrichment of functions related to chemotaxis and production of chemicals with potential antimicrobial properties, including terpenoids and polyketides, in thermosensitive corals suggests the presence and activity of opportunistic bacteria (Raimundo et al., 2018). Chemotaxis is associated with virulence of pathogenic bacteria and flagellar proteins are abundant in known pathogens like *Vibrio shilonii* (Li et al., 2016). *Vibrio coralliilyticus* also uses chemotaxis elicited by increased DMSP concentrations to locate the mucus of its heat-stressed host (Gao et al., 2021; Garren et al., 2013) and its flagellar proteins have been shown to be important for adhesion to the coral during infection (Akahoshi and Bevins, 2022; Ramos et al., 2004; Meron et al., 2009). Terpenoids produced by microorganisms are associated with microbial competition, defense and quorum sensing for opportunistically pathogenic microbes and can play a role in their adaptation to common stressors (Avalos et al., 2021). Bacteria have also been observed to use polyketides to defeat other microbial competitors and to enable pathogenesis by suppressing the immune response of the host (Ridley et al, 2007). It is important to note, however, that the PICRUSt2 method that was used to derive microbiome functions is purely predictive. Nevertheless, it provides a starting point for understanding gene functions potentially represented within a microbial community. Metagenome or metatranscriptome studies would be critical to provide further insights into functions that may differ in microbiomes associated with coral tolerance or susceptibility to thermal stress.

### Harnessing the microbiome to promote thermotolerance

Microbiome responses in the coral holobiont are driven not only by interactions within the microbial community, but also by interactions with the host. The coral regulates its microbiome through physical barriers such as the mucus layer (Ritchie, 2006), through manipulation of symbiont physiology (Voolstra et al., 2024; Wilde et al., 2024), and through innate immune functions that mediate symbiont recognition, maintenance and dysbiosis (Weis, 2008; Posadas et al., 2021; Mohamed et al., 2023). The coral host can distinguish beneficial versus harmful microbes through pattern recognition receptors (PRRs) on phagocyte cell surfaces that recognize microbial-associated molecular patterns (MAMPs) on microbial cell membranes (Weis, 2008). Although we did not determine host genotypes in this study, it is possible that a disparate repertoire of PRRs may contribute to microbial community differences associated with coral thermotolerance categories. Thus, further investigations are needed to assess host genotype-phenotype connection to microbiome structure (Glasl et al., 2019), and to elucidate the mechanisms regulating microbiome structure in thermotolerant and thermosensitive coral hosts.

With increasing interest in harnessing beneficial microorganisms for corals (BMCs) to enhance coral tolerance and mitigate temperature stress (Rosado et al., 2018; Epstein et al., 2019; Santoro et al., 2021; Doering et al., 2023), understanding the factors that drive coral microbial community structure is crucial. Studies that have tested the efficacy of BMCs on the performance of corals under thermal stress have so far been promising (Peixoto et al., 2017; Santoro et al., 2021; Li et al., 2022; Delgadillo-Ordoñez et al., 2024). However, a major challenge for this approach is that inoculated microbes are not stably integrated or retained by the recipient colony, likely due to a combination of microbiome and host-mediated regulation (Rosado et al., 2019; Santoro et al., 2021; Zhang et al., 2021; Delgadillo-Ordoñez et al., 2024). By comparing microbiomes of thermotolerant and thermosensitive corals, we identified potential BMCs that are ubiquitous and present at greater abundance in thermotolerant colonies. We infer that supplementing thermosensitive colonies with these ‘native’ beneficial bacteria may improve the efficacy of this approach.

Some potential BMCs in the *A.* cf. *tenuis* microbiome include *Endozoicomonas*, *Pseudoaltermonas*, and *Bdellovibrio*. Inoculation of *Pseudoalteromonas* strains and *Halomonas* have been reported to mitigate coral bleaching due to thermal and pathogen stresses in *P. damicornis* (Rosado et al., 2019). Consistent with this, transplantation of the *Endozoicomonas*- dominated microbiome of heat-tolerant *Porites* sp. and the predatory *Bdellovibrio*-containing microbiome of heat-tolerant *Pocillopora* into heat-sensitive conspecific recipients alleviated bleaching from short-term thermal stress exposure (Doering et al., 2021). Culturing of BMCs from heat-tolerant *A*. cf. *tenuis* will enable future experiments to test their ability to boost thermotolerance of sensitive colonies.

## CONCLUSION

The results of this study demonstrate that microbial community structure does not directly correlate with coral thermotolerance under ambient temperature conditions. Microbiome responses to thermal stress varied, with only thermosensitive colonies exhibiting a marked shift in community structure that coincided with bleaching. The presence of specific microbial taxa in the coral community may serve as a predictor of performance under stress. Thermotolerant colonies contained diverse and abundant microbes belonging to potentially beneficial taxa such as *Endozoicomonas*, *Arcobacter*, *Bifidobacterium* and *Lactobacillus*. Thermosensitive colonies, on the other hand, exhibited depletion of these potentially beneficial microbes and an increase in opportunistic pathogens (e.g. Flavobacteriales, Rhodobacteraceae, *Vibrio*) when subjected to elevated temperature. This information on microbial signatures associated with coral thermotolerance may be harnessed for applications that aim to increase the vigor of reef-building corals and maintain reef health through a healthy microbiome.

## AUTHOR CONTRIBUTIONS

**Jake Ivan P. Baquiran:** Conceptualization; data curation; formal analysis; investigation; methodology; writing – original draft preparation; writing – review & editing. **John Bennedick Quijano:** Data curation; formal analysis; methodology; software; visualization; writing – original draft preparation; writing – review & editing. **Madeleine J.H. van Oppen:** Funding acquisition; writing – original draft preparation; writing – review & editing. **Patrick C. Cabaitan:** Funding acquisition; writing – original draft preparation; writing – review & editing. **Peter L. Harrison:** Funding acquisition; writing – original draft preparation; writing – review & editing. **Cecilia Conaco:** Conceptualization; funding acquisition; methodology; supervision; writing – original draft preparation; writing – review & editing.

## ACKNOWLEDGEMENTS

The authors greatly thank Ben Jack Gabuay and Rickdane Gomez for their invaluable assistance in the field surveys, coral collection, and experimentation. This study was funded by the Department of Science and Technology Philippine Council for Agriculture, Aquatic and Natural Resources Research and Development (QMSR- MRRD-MEC-295-1449) to PCC and CC and by an Australian Centre for International Agricultural Research (ACIAR) grant FIS/2019/123 to PLH, MJHvO, PCC, and CC.

