## Supplementary Materials for "Microbiome stability is linked to coral thermotolerance"

Running title: Heat tolerant corals have stable microbiome

Jake Ivan P. Baquiran[1,2†], John Bennedick Quijano[1†], Madeleine J.H. van Oppen[3,4], Patrick C. Cabaitan[1], Peter L. Harrison[3,5] and Cecilia Conaco[1]*

[1]Marine Science Institute, University of the Philippines Diliman, Quezon City, 1101, Philippines
[2]Graduate School of Engineering and Science, University of the Ryukyus, 1 Senbaru, Nishihara, Okinawa 903-0213, Japan
[3]Australian Institute of Marine Science, PMB No 3, Townsville MC, Queensland 4810, Australia
[4]School of BioSciences, University of Melbourne, Parkville, Victoria 3010, Australia
[5]Faculty of Science and Engineering, Southern Cross University, Lismore, NSW 2480, Australia

*Correspondence:
Cecilia Conaco


†These authors contributed equally to this work and share first authorship

**Supplementary Figures**

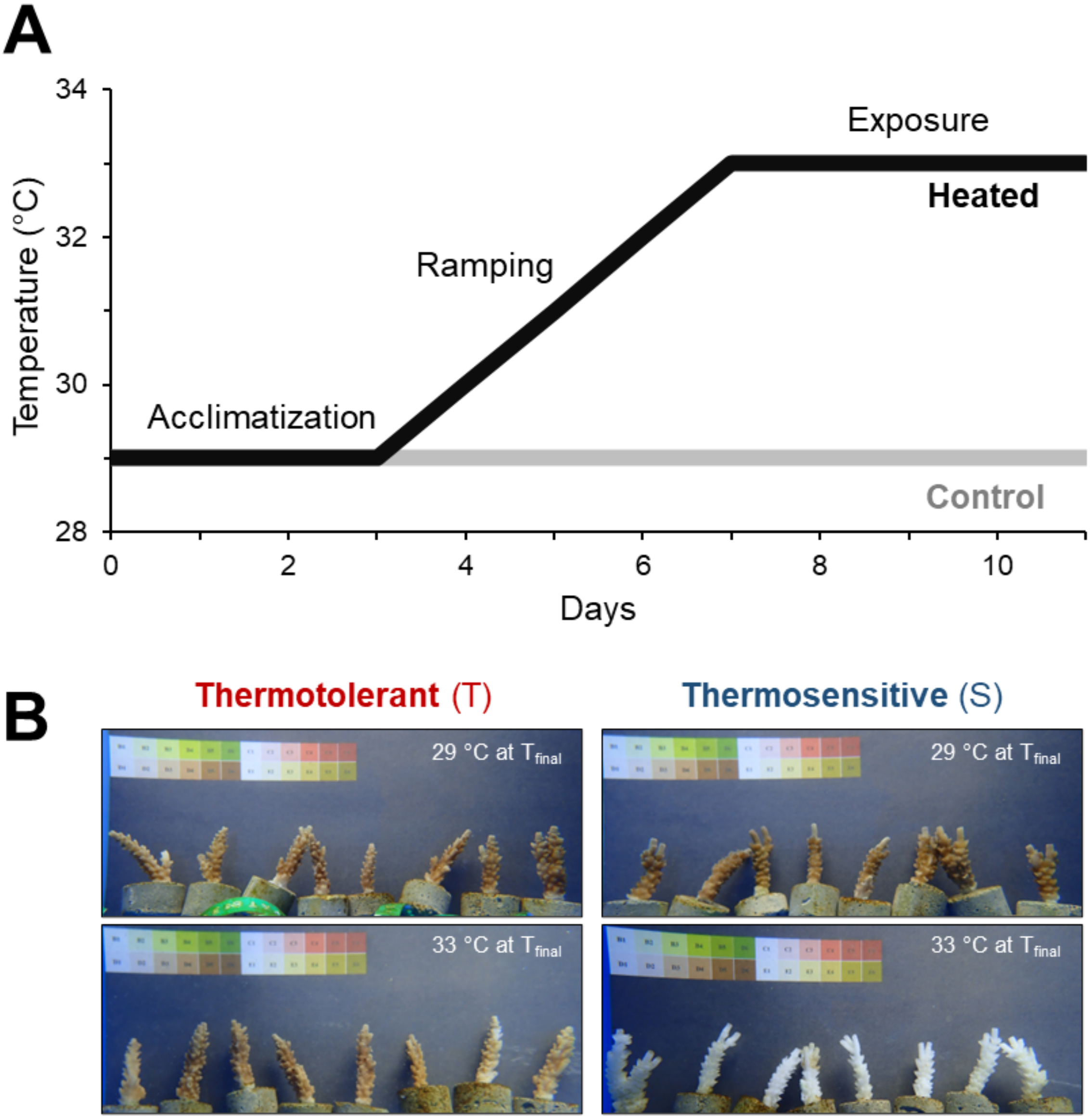


**Supplementary Figure 1.** Heat tolerance variation in *A.* cf. *tenuis* colonies. (a) Temperature profile for the short-term thermal stress experiment. (b) Representative images of fragments from thermotolerant (colony 3) and thermosensitive (colony 1) *A.* cf. *tenuis* colonies after 4 days sustained exposure to 29 or 33°C ($T_{final}$).

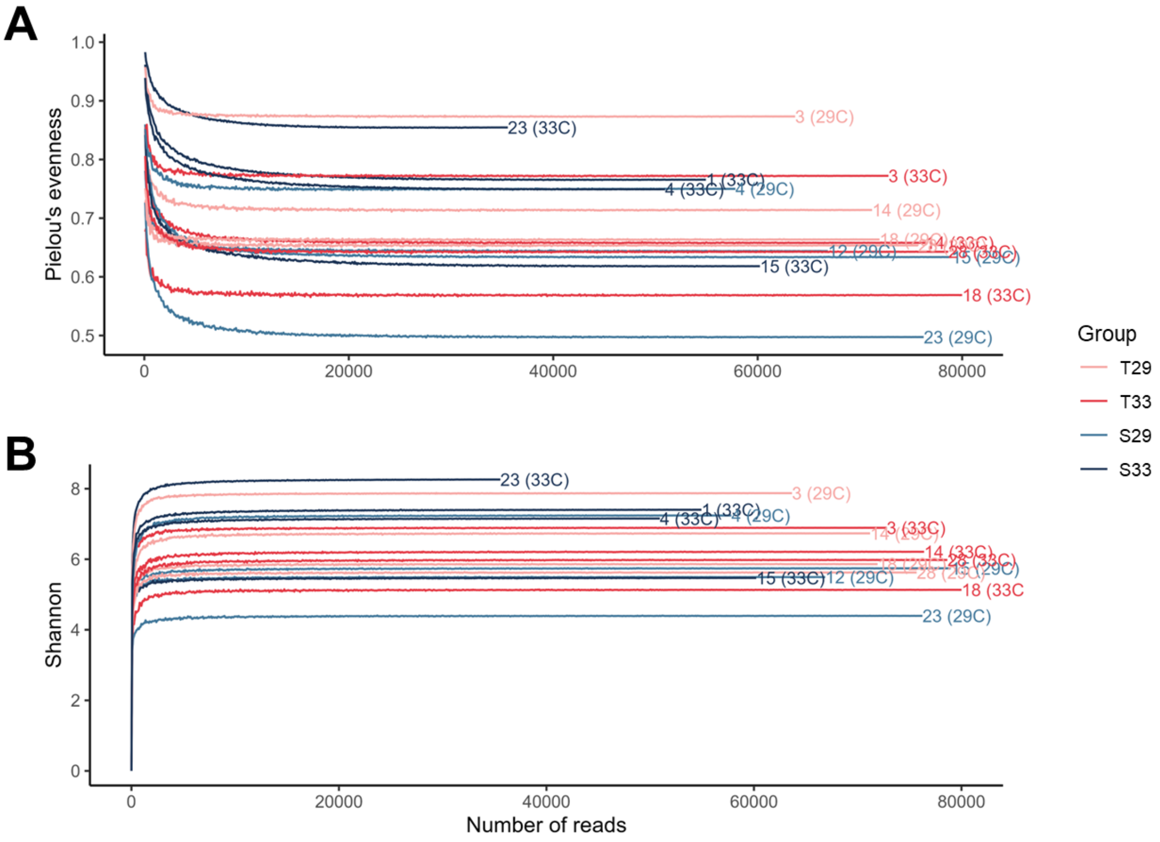


**Supplementary Figure 2.** Rarefaction curves based on (a) Pielou's evenness and (b) Shannon index.

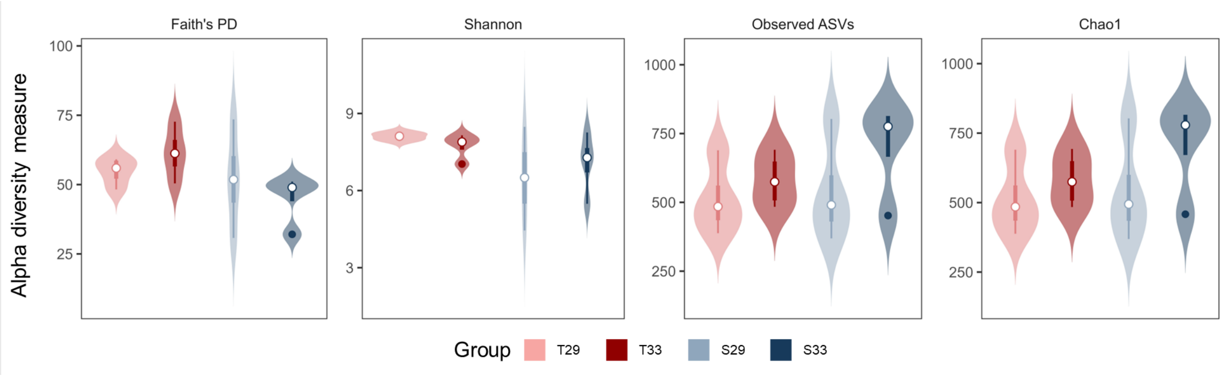


**Supplementary Figure 3.** Alpha diversity measures compared among heat tolerance classifications and temperature treatments. No significant differences were observed for all the measures among groups (Kruskal-Wallis, p-value >0.05).

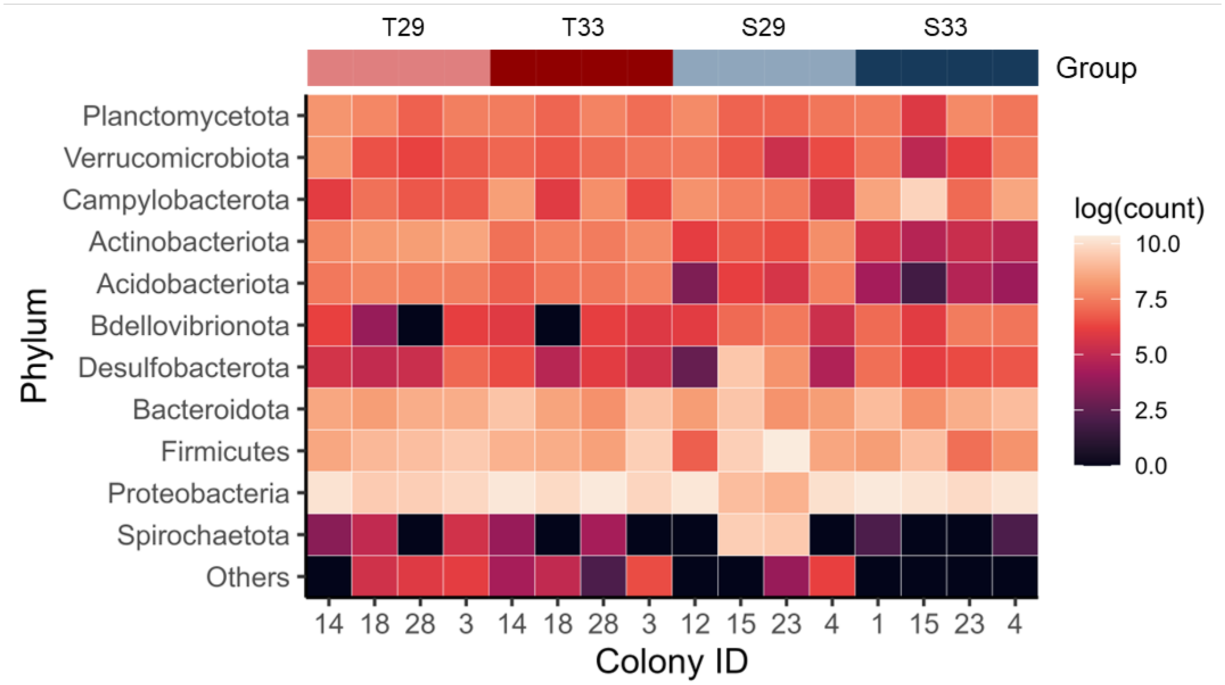


**Supplementary Figure 4.** Relative abundance of major (>1% total relative abundance) microbial phyla in the *A.* cf. *tenuis* microbiome.

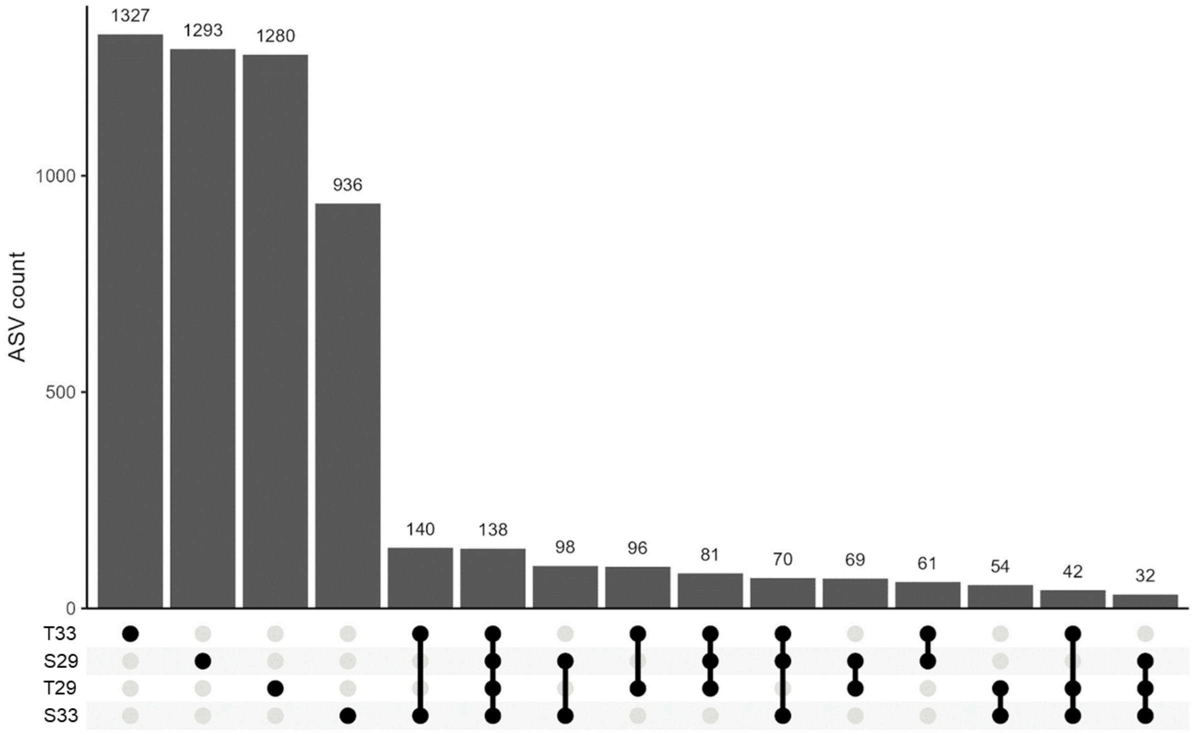


**Supplementary Figure 5.** Number of shared and unique ASVs among corals from different heat tolerance classifications and treatment groups.

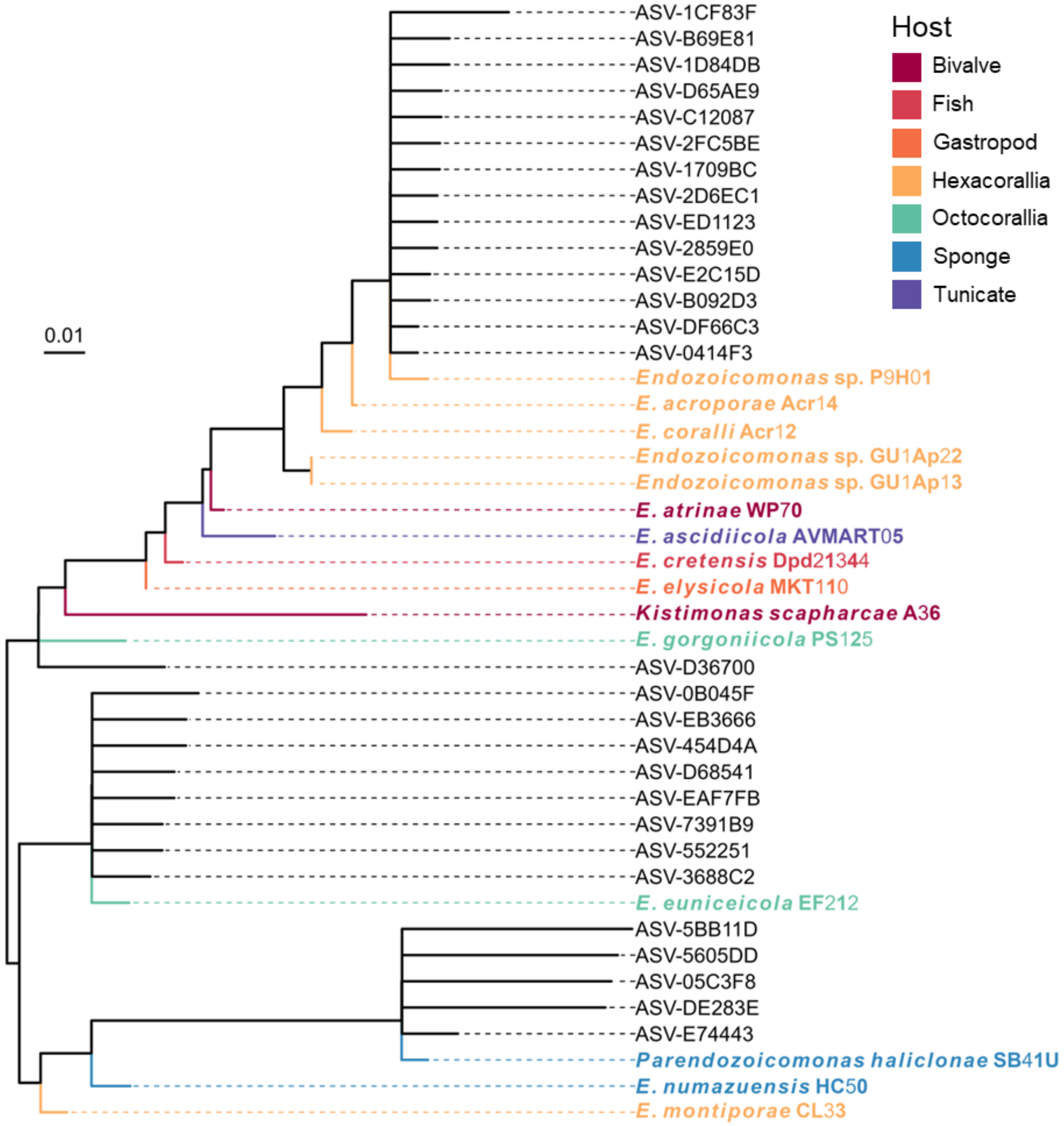


**Supplementary Figure 6.** Phylogenetic placement of *Endozoicomonas* ASVs from *A.* cf. *tenuis* on an ML tree based on near full-length *Endozoicomonas* 16S rRNA sequences using EPA-NG (Barbera et al., 2018). Names in bold represent reference sequences. Colors represent the host group from which the isolates originated.

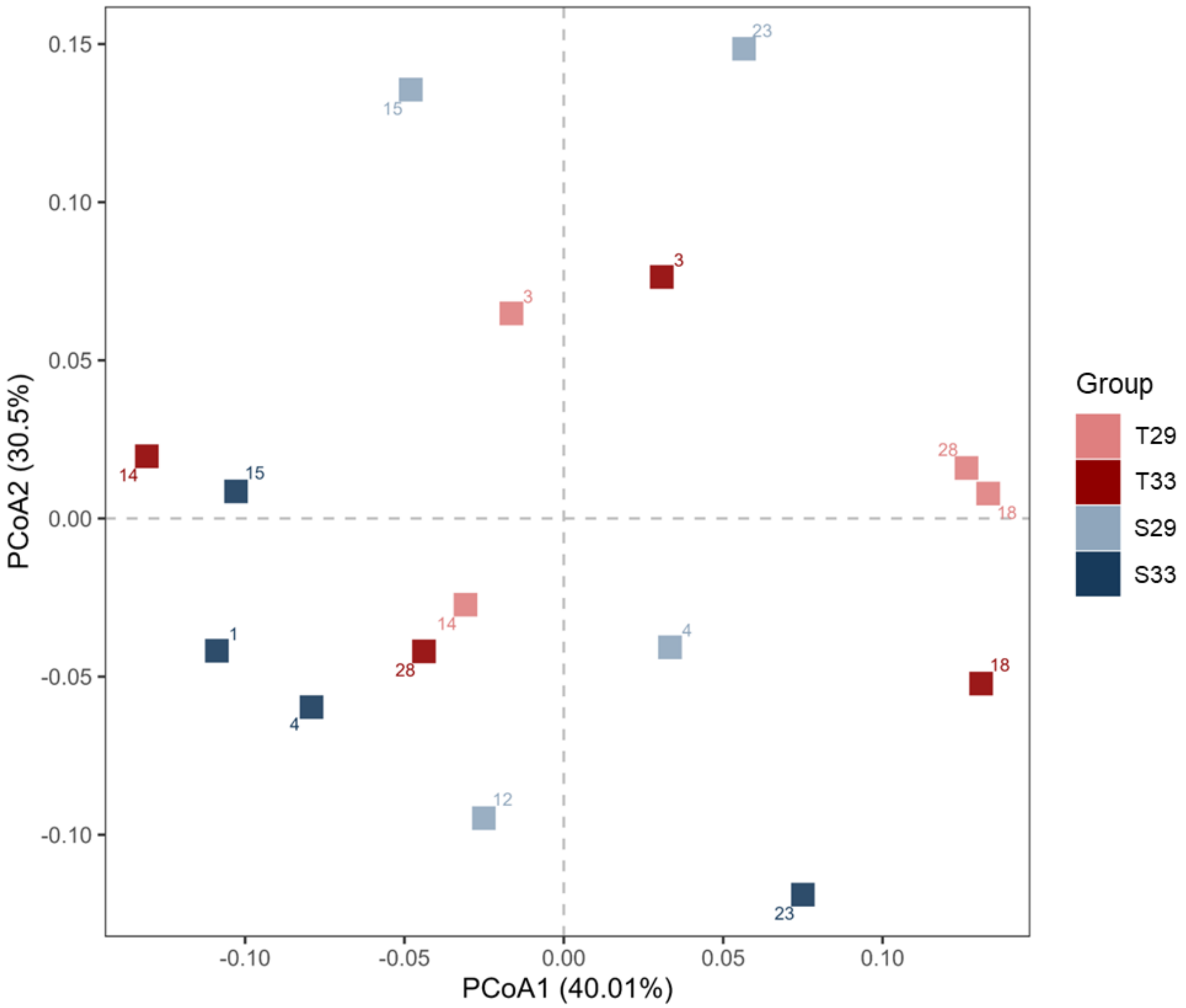


**Supplementary Figure 7.** PCoA plot of predicted functions of the *A.* cf. *tenuis* microbiome. Plot is based on Bray-Curtis dissimilarity of PICRUSt2-predicted functions. Colony IDs are shown beside the markers.

**Supplementary Tables**

**Supplementary Table 1.** Physicochemical parameters in control and heated tanks during the sustained exposure stage of the thermal stress experiment (days 8 to 11). Values represent mean ± standard error.

| Treatment | Temperature (°C) | Salinity (ppt) | Dissolved oxygen (mg/L) | pH |
|---|---|---|---|---|
| 29°C | 28.77±0.02 | 32.85±0.11 | 8.14±0.05 | 8.11±0.01 |
| 33°C | 33.12±0.02 | 31.93±0.06 | 7.86±0.05 | 8.09±0.01 |

**Supplementary Table 2.** Bleaching proportion of fragments after 4 days of sustained exposure to 33°C.

| Colony ID | Category | No bleaching (%) | Partial bleaching (%) | Complete bleaching (%) |
|---|---|---|---|---|
| 3 | T | 100 | 0 | 0 |
| 14 | T | 100 | 0 | 0 |
| 18 | T | 100 | 0 | 0 |
| 28 | T | 100 | 0 | 0 |
| 1 | S | 0 | 0 | 100 |
| 4 | S | 0 | 0 | 100 |
| 12 | S | 50 | 0 | 50 |
| 15 | S | 12.5 | 12.5 | 75 |
| 23 | S | 0 | 0 | 100 |

**Supplementary Table 3.** Comparison of dispersion or variability of *Acropora* cf. *tenuis* microbial communities at ASV level among tolerance classifications and temperature treatments based on Bray-Curtis dissimilarity.

| Group | Parameters | Df | Sum Sq | Mean Sq | F value | Pr(>F) |
|---|---|---|---|---|---|---|
| Tolerance (T or S) | Groups | 1 | 0.0034 | 0.0034 | 0.3288 | 0.5755 |
| | Residuals | 14 | 0.1433 | 0.0102 | - | - |
| Treatment (29 or 33°C) | Groups | 1 | 0.0160 | 0.0160 | 1.6808 | 0.2158 |
| | Residuals | 14 | 0.1337 | 0.0095 | - | - |
| Tolerance x treatment (S29, T29, S33, T33) | Groups | 3 | 0.1114 | 0.0371 | 3.1887 | 0.0628 |
| | Residuals | 12 | 0.1397 | 0.0116 | - | - |

**Supplementary Table 4.** List of differentially abundant microbial ASVs based on ANCOMBC analysis (p-value < 0.05). Positive log-fold change indicates enrichment and negative log-fold change indicates depletion.

| ASV | Log-fold change | Phylum | Class | Order | Family | Genus | Most resolved taxon |
|---|---|---|---|---|---|---|---|
| ASV-F70375 | 2.23508923 | Proteobacteria | Gammaproteobacteria | Arenicellales | Arenicellaceae | NA | Arenicellaceae |
| ASV-9BA24A | 2.3608036 | Proteobacteria | Gammaproteobacteria | Pseudomonadales | Cellvibrionaceae | NA | Cellvibrionaceae |
| ASV-CE0D4E | 3.39554839 | Proteobacteria | Gammaproteobacteria | Pseudomonadales | Spongiibacteraceae | NA | Spongiibacteraceae |
| ASV-32EBDF | 5.00628125 | Proteobacteria | Gammaproteobacteria | Pseudomonadales | NA | NA | Pseudomonadales |
| ASV-DF3006 | 2.80750469 | Bacteroidota | Bacteroidia | Flavobacteriales | Cryomorphaceae | Owenweeksia | Owenweeksia |
| ASV-9FC722 | 3.0410358 | Bacteroidota | Bacteroidia | Flavobacteriales | Flavobacteriaceae | Pseudofulvibacter | Pseudofulvibacter |
| ASV-5342E0 | 2.7962958 | Bacteroidota | Bacteroidia | Chitinophagales | NA | NA | Chitinophagales |
| ASV-43392C | 2.20991168 | Proteobacteria | Gammaproteobacteria | Pseudomonadales | Saccharospirillaceae | NA | Saccharospirillaceae |
| ASV-DFCF4A | 3.90768411 | Proteobacteria | Gammaproteobacteria | Enterobacterales | Pseudoalteromonadaceae | Pseudoalteromonas | Pseudoalteromonas |
| ASV-35DAB5 | 3.05282279 | Proteobacteria | Alphaproteobacteria | Rhodobacterales | Rhodobacteraceae | NA | Rhodobacteraceae |
| ASV-487CBA | 3.55496297 | Proteobacteria | Alphaproteobacteria | NA | NA | NA | Alphaproteobacteria |
| ASV-F64B0D | 3.65121565 | Proteobacteria | Gammaproteobacteria | Pseudomonadales | Saccharospirillaceae | NA | Saccharospirillaceae |
| ASV-29420F | 3.00101485 | Bacteroidota | Bacteroidia | Flavobacteriales | Flavobacteriaceae | NA | Flavobacteriaceae |
| ASV-DBAE81 | 2.63351162 | Bacteroidota | Bacteroidia | Flavobacteriales | NA | NA | Flavobacteriales |
| ASV-5FC884 | 3.16974947 | Bacteroidota | Bacteroidia | Flavobacteriales | Crocinitomicaceae | Crocinitomix | Crocinitomix |
| ASV-A44B39 | 4.31051395 | Myxococcota | Polyangia | NA | NA | NA | Polyangia |
| ASV-E0E1AB | 3.37281016 | Planctomycetota | Phycisphaerae | Phycisphaerales | Phycisphaeraceae | NA | Phycisphaeraceae |
| ASV-15C525 | 3.10490164 | Desulfobacterota | Desulfuromonadia | Bradymonadales | Bradymonadales | Bradymonadales | Bradymonadales |
| ASV-592A98 | 1.96600713 | Bdellovibrionota | Bdellovibrionia | Bacteriovoracales | Bacteriovoraceae | NA | Bacteriovoraceae |
| ASV-012BF8 | 4.28739577 | Bacteroidota | Bacteroidia | Chitinophagales | Saprospiraceae | NA | Saprospiraceae |
| ASV-45A7E1 | 3.35450093 | Bdellovibrionota | Bdellovibrionia | Bdellovibrionales | Bdellovibrionaceae | NA | Bdellovibrionaceae |
| ASV-D6DD49 | 2.83725096 | Proteobacteria | Gammaproteobacteria | Pseudomonadales | Oleiphilaceae | Oleiphilus | Oleiphilus |
| ASV-4CFB3C | 3.78539137 | Proteobacteria | Gammaproteobacteria | Enterobacterales | Vibrionaceae | Vibrio | Vibrio |
| ASV-5C73EC | 3.16480954 | Bacteroidota | Bacteroidia | Flavobacteriales | Crocinitomicaceae | Fluviicola | Fluviicola |
| ASV-4CC4ED | 3.27669401 | Bacteroidota | Bacteroidia | Flavobacteriales | NA | NA | Flavobacteriales |
| ASV-611E18 | 3.54044081 | Proteobacteria | Gammaproteobacteria | Pseudomonadales | Nitrincolaceae | Nitrincolaceae | Nitrincolaceae |
| ASV-54F506 | 3.71134881 | Bacteroidota | Bacteroidia | Flavobacteriales | Crocinitomicaceae | Crocinitomix | Crocinitomix |
| ASV-DB5E82 | 3.33115772 | Bdellovibrionota | Bdellovibrionia | Bdellovibrionales | Bdellovibrionaceae | NA | Bdellovibrionaceae |
| ASV-F9266D | 2.72819727 | Desulfobacterota | Desulfuromonadia | Bradymonadales | Bradymonadales | Bradymonadales | Bradymonadales |
| ASV-B90ACA | 3.34931115 | Myxococcota | Polyangia | NA | NA | NA | Polyangia |

| | | | | | | | |
|---|---|---|---|---|---|---|---|
| ASV-60AD81 | 2.61305363 | Bdellovibrionota | Bdellovibrionia | Bdellovibrionales | Bdellovibrionaceae | NA | Bdellovibrionaceae |
| ASV-C2FD58 | 2.27912361 | Bacteroidota | Bacteroidia | Flavobacteriales | Cryomorphaceae | NA | Cryomorphaceae |
| ASV-04C081 | 4.62884409 | Bacteroidota | Bacteroidia | Flavobacteriales | Flavobacteriaceae | Tenacibaculum | Tenacibaculum |
| ASV-1582AF | 3.22076522 | Proteobacteria | Alphaproteobacteria | Rhodospirillales | Terasakiellaceae | NA | Terasakiellaceae |
| ASV-C6C054 | 3.36534833 | Bacteroidota | Bacteroidia | Flavobacteriales | Flavobacteriaceae | Tenacibaculum | Tenacibaculum |
| ASV-7793E4 | 2.19828167 | Bacteroidota | Bacteroidia | Chitinophagales | Saprospiraceae | NA | Saprospiraceae |
| ASV-D25BEE | 3.72097473 | Proteobacteria | Gammaproteobacteria | Enterobacterales | Shewanellaceae | Ferrimonas | Ferrimonas |
| ASV-A81B3A | 3.13307266 | Proteobacteria | Gammaproteobacteria | Pseudomonadales | NA | NA | Pseudomonadales |
| ASV-D8F9B1 | 3.47796737 | Proteobacteria | Gammaproteobacteria | Pseudomonadales | NA | NA | Pseudomonadales |
| ASV-92FEA1 | 2.84703101 | Proteobacteria | Gammaproteobacteria | Pseudomonadales | NA | NA | Pseudomonadales |
| ASV-D8F12F | 3.08201774 | Proteobacteria | Gammaproteobacteria | Burkholderiales | Methylophilaceae | Methylotenera | Methylotenera |
| ASV-B7BB8A | 3.38496497 | Bacteroidota | Bacteroidia | Flavobacteriales | Flavobacteriaceae | Gilvibacter | Gilvibacter |
| ASV-12CAFC | 2.72733071 | Actinobacteriota | Acidimicrobiia | Microtrichales | Ilumatobacteraceae | NA | Ilumatobacteraceae |
| ASV-118FCA | 5.90117199 | Campylobacterota | Campylobacteria | Campylobacterales | Arcobacteraceae | Malaciobacter | Malaciobacter |
| ASV-28B9EF | 2.10539078 | Bacteroidota | Bacteroidia | Flavobacteriales | Cryomorphaceae | NA | Cryomorphaceae |
| ASV-2FE7AD | 3.05355046 | Proteobacteria | Alphaproteobacteria | Rhodobacterales | Rhodobacteraceae | NA | Rhodobacteraceae |
| ASV-11AFDA | 3.04663035 | Desulfobacterota | Desulfuromonadia | NA | NA | NA | Desulfuromonadia |
| ASV-5A9A01 | 2.23236397 | Bacteroidota | Bacteroidia | Flavobacteriales | Flavobacteriaceae | Tenacibaculum | Tenacibaculum |
| ASV-F58DA0 | 3.02903515 | Proteobacteria | Gammaproteobacteria | Pseudomonadales | NA | NA | Pseudomonadales |
| ASV-7391B9 | -4.9817507 | Proteobacteria | Gammaproteobacteria | Pseudomonadales | Endozoicomonadaceae | Endozoicomonas | Endozoicomonas |
| ASV-2753C5 | -3.8534186 | Synergistota | Synergistia | Synergistales | Synergistaceae | Pyramidobacter | Pyramidobacter |

**Supplementary Table 5.** Comparison of *Endozoicomonas* ASV abundance in the *Acropora* cf. *tenuis* microbial community for different tolerance classifications and temperature treatments based on Bray-Curtis dissimilarity. Benjamini-Hochberg adjusted p-values in bold denote statistical significance at $p < 0.05$.

| **Comparison** | **PERMANOVA** | |
|---|---|---|
| | **pseudo-F** | **p-value** |
| T29 x T33 | 0.7619 | 0.639 |
| T29 x S29 | 0.535 | 0.883 |
| T29 x S33 | **3.014** | **0.048** |
| T33 x S29 | 0.861 | 0.496 |
| T33 x S33 | **3.4327** | **0.034** |
| S29 x S33 | 2.1476 | 0.188 |

**Data Files**

**Data File 1.** Abundance table at ASV level with taxon affiliations
**Data File 2.** List of differentially abundant ASVs based on ANCOMBC analysis (p-value < 0.05)
**Data File 3.** Differentially abundant predicted microbial functions based on KEGG ortholog (KO) genes BRITE Level 2
